# Aberrant cortical–subcortical somatosensory network organization is associated with impaired tactile-motion perception in akinetic-rigid Parkinson’s disease

**DOI:** 10.64898/2026.05.21.726534

**Authors:** Boyi Qu, Yasi Jiang, Yumei Yue, Ting Shen, Chaohong Gao, Dian Huang, Ying Shen, Chris De Zeeuw, Baorong Zhang, Hsin-Yi Lai

**Author notes:** **Correspondence:** Hsin-Yi Lai,; Baorong Zhang.

## Abstract

**Background:** Parkinson’s disease (PD) is associated with somatosensory dysfunction, yet higher-order tactile-motion processing and its neural basis remain poorly understood. Here, we combined psychophysical assessment of tactile-motion perception with resting-state functional MRI (rs-fMRI) to investigate how altered sensorimotor network organization relates to tactile perceptual deficits in akinetic-rigid dominant PD (PD_AR_).

**Methods:** Forty-six participants (25 PD_AR_ and 21 healthy controls, HCs) underwent clinical evaluation and resting-state functional MRI scanning. A subset of participants (15 PD_AR_ and 15 HCs) additionally completed a tactile-motion perception task assessing directional bias and speed sensitivity. ROI-to-ROI functional connectivity analyses were performed within a customized 26-node sensorimotor and cortico-basal ganglia-thalamo-cortical (CBGTC) network.

**Results:** Compared with HCs, PD_AR_ patients exhibited significantly altered tactile-motion perception, including greater directional bias (*p* = 0.047) and reduced sensitivity to dynamic tactile speed (*p* < 0.001), reflecting diminished sensitivity to mid-to-high speeds. Resting-state functional connectivity analyses revealed widespread reductions in connectivity within cortical somatosensory regions (Areas 1, 2, 3a, 3b, 5), particularly between primary somatosensory and motor cortices. In contrast, increased connectivity was observed between basal ganglia-thalamic structures and cortical sensorimotor regions, including thalamus–M1 and GPi–sensorimotor connections (all *p*-FDR < 0.05).

**Conclusions:** These findings demonstrate a dual pattern of sensorimotor network reorganization in PD_AR_, characterized by disrupted cortical somatosensory integration alongside enhanced subcortical-cortical coupling. The present study provides evidence that higher-order tactile-motion dysfunction in PD is associated with large-scale abnormalities in distributed sensorimotor networks and highlights the somatosensory system as a potential target for future therapeutic intervention.

## 1 Introduction

Parkinson’s disease (PD) is a progressive neurodegenerative disorder characterized by hallmark motor symptoms such as tremor, rigidity, and bradykinesia, alongside a spectrum of non-motor manifestations that often precede or exacerbate motor impairments (Schapira et al. 2017). Among these, somatosensory dysfunction, including deficits in tactile perception and proprioception, has emerged as a clinically significant yet underexplored domain (Juri et al. 2010; Nolano et al. 2008). Such abnormalities disrupt sensorimotor integration, leading to impaired motor planning and execution, excessive grip force (Davidson et al. 2025), reduced spatial acuity (Konczak et al. 2012), and diminished responsiveness to dynamic tactile inputs during daily activities (Foki et al. 2015; Lee et al. 2018). At the neural level, dopamine depletion is thought to perturb somatosensory processing through large-scale disturbances within the striato-thalamo-cortical (STC) loop (Nagy et al. 2006), weakening sensory relay and broadening receptive fields, thereby introducing neural noise and elevating perceptual thresholds (A. Conte et al. 2013; Obeso et al. 2008). These central alterations impair sensory integration and contribute to impaired sensorimotor coordination underlying parkinsonian motor dysfunction.

Resting-state functional MRI (rs-fMRI) provides a powerful approach to characterize intrinsic neural alterations in PD without task demands (Tessitore et al. 2019). Accumulating evidence has revealed widespread disruptions in functional connectivity across large-scale networks, particularly within the sensorimotor network (SMN). Altered neural activity and connectivity have been observed not only in primary motor and somatosensory cortices but also in associative cortical regions involved in higher-order integration (Permezel et al. 2023; S. Wang et al. 2021; Y. Wang et al. 2024). Notably, the primary somatosensory cortex (S1), a key hub for tactile and proprioceptive processing, exhibits marked connectivity alterations in PD. Advanced analyses have further identified disrupted feedforward sensorimotor pathways and abnormal temporal connectivity states characterized by global hypo-connectivity alongside potential compensatory cerebellar-motor interactions (Goelman et al. 2021; Shen et al. 2024). Despite these insights, it remains unclear whether higher-order tactile processing deficits in PD arise primarily from localized somatosensory dysfunction or from broader network-level dysfunction across cortical and subcortical sensorimotor circuits.

While behavioral studies have reported somatosensory impairments in PD, most have focused on simple detection or vibration tasks, leaving higher-order processes such as tactile-motion perception largely unexplored. Previous findings indicate that PD patients exhibit reduced accuracy and increased perceptual bias and prolonged response times during basic tactile detection tasks (Antonella Conte et al. 2017; Fiorio et al. 2014; Kesayan et al. 2015). However, it remains unclear whether these deficits arise from localized somatosensory dysfunction or from broader network-level disturbances spanning cortical and subcortical sensorimotor circuits. Because tactile-motion perception depends critically on the integration of spatial and temporal sensory information, combining psychophysical assessment with rs-fMRI enables direct investigation of how disrupted cortical integration and abnormal subcortical coupling jointly shape tactile-motion perception in PD.

To address this gap, we combined psychophysical assessment of tactile-motion perception with rs-fMRI in individuals with PD and age-matched healthy controls. This multimodal framework allowed us to examine how disruptions in intrinsic functional connectivity within somatosensory cortical regions and cortico-basal ganglia-thalamo-cortical (CBGTC) contribute to tactile-motion perceptual deficits in PD. We hypothesized that PD patients would exhibit deficits in tactile-motion perception accompanied by a network-level imbalance characterized by cortical sensory disintegration and abnormal cortico-subcortical connectivity alterations. Elucidating these associations may provide mechanistic insight into how disrupted sensorimotor network organization contributes to tactile dysfunction and motor impairment in PD.

## 2 Methods

### 2.1 Subjects

This study was approved by the Human Research Ethics Committee of the Second Affiliated Hospital, Zhejiang University School of Medicine (IRB No.2016-002). Twenty-five patients with PD (12 males; mean age = 57.1±7.2 years) and 21 age- and sex-matched healthy controls (HCs; 11 males; mean age = 53.9±5.0 years) were recruited from the Department of Neurology of Second Affiliated Hospital and Hangzhou community. All participants were right-handed, as assessed by the Edinburgh Handedness Inventory (Oldfield 1971).

The diagnosis of PD was confirmed by experienced neurologists according to the Movement Disorder Society (MDS) Clinical Diagnostic Criteria for Parkinson’s disease (Postuma et al. 2015). Each patient underwent comprehensive neurological assessments, including the MDS-sponsored revision of the Unified Parkinson’s Disease Rating Scale (MDS-UPDRS), Hoehn & Yahr (H-Y) Scale, Mini-mental State Examination (MMSE), and Montreal Cognitive Assessment (MoCA). Exclusion criteria were atypical parkinsonism, other neurologic or psychiatric disorders, history of stroke or dementia, use of neuroleptic medication, and general contraindications for MRI.

All patients were classified as akinetic-rigid dominant PD (PD_AR_) according to the modified UPDRS-III ratio proposed by Schiess’s group (Schiess et al. 2000), with a ratio < 1.0. Neurological examinations, MRI scanning and tactile-motion perception experiments were conducted in the practically defined “off-medication” state. All PD_AR_ patients and HCs underwent MRI scanning. A subset of participants, including 15 PD_AR_ patients (9 males, 55.5 ± 4.9 years) and 15 HCs (7 males, 52.9 ± 6.5 years), also completed the tactile-motion perception experiment.

### 2.2 Tactile-motion perception experiment and data analysis

To assess the tactile-motion perception in PD_AR_ patients, we employed a custom-built tactile stimulator (Figure 1A) capable of delivering precise motion stimuli to the distal fingerpad (Qu et al. 2025). The device consisted of three miniature motors (1024M006S, 1516E006SR and AM1524, Faulhaber, Switzerland) that controlled the rotation and indentation of a 20-mm aluminum stimulus ball engraved with 4-mm square-wave grating. Participants were seated comfortably with the left forearm fixed in a vertical position and the index finger (D2) gently in contact with the stimulus ball. A touchpad (PTH-660, Wacom Co. Ltd., Japan) was placed between the participants’ eyes and the stimulator to block visual feedback, while continuous white noise was delivered through headphones to mask motor sounds. Each trial presented a tactile stimulus with 0.5-mm indentation, eight directions (0° to 157.5°, 22.5° steps), and six speeds (20, 40, 80, 160, 240, 320 mm/s), each lasting 1 s.

**Figure 1.**
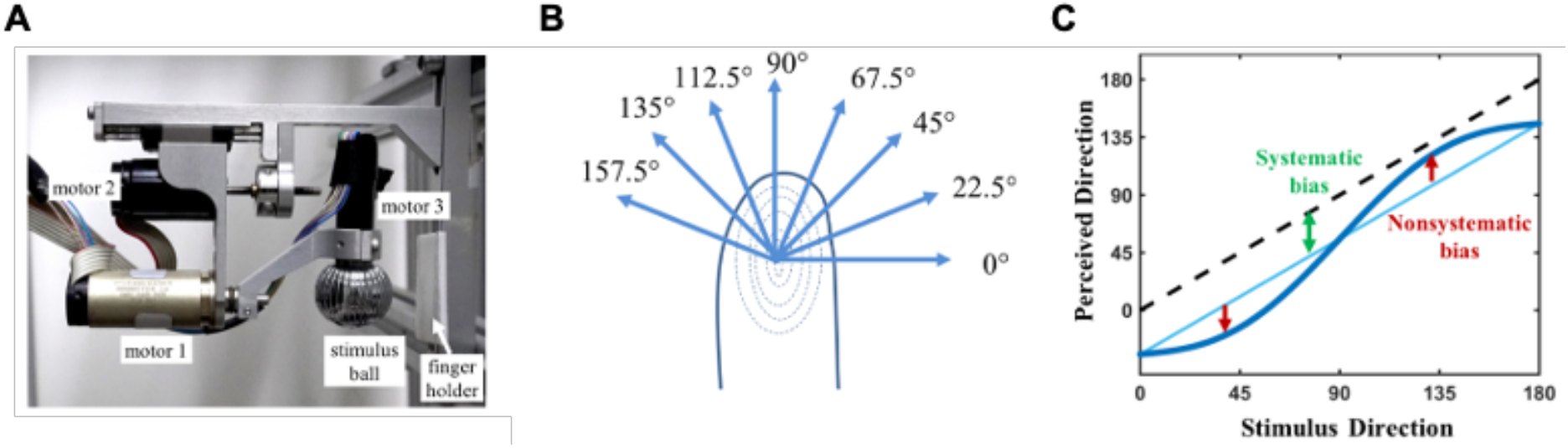
A setup of experiment. (A) Stimuli apparatus of tactile stimulator. Motor 1, motor 2 and motor 3 were used to control the direction of stimulus ball, depth of indentation to the skin, and control the rotation speed of stimulus ball. (B) The stimulus parameters included 8 stimulus directions with 22.5-degree step. (C) The perceptual bias of direction including the systematic bias and the nonsystematic bias was fitted by linear model and cosine function, respectively. The green arrow represented the systematic bias and the red arrows represented the nonsystematic bias.

A total of 240 trials (8 directions x 6 speeds x 5 repetitions) were administered. Participants rested for 3 mins between repetitions to minimize fatigue and attentional drift. After each trial, participants reported the perceived direction and speed of tactile motion using the right index finger on the touchpad. Response trajectories were recorded at a sampling rate of 200 Hz and calculated by MATLAB (R2018b, MathWorks, Natick, MA).

The perceived direction and speed of stimulus were analyzed using different fitting models (Chen et al. 2020). The difference between the reported direction (*RD*_*i*_) and stimulus direction (*SD*_*i*_) was defined as the perceptual bias of direction (*Pb*_*i*_) calculating as follow:

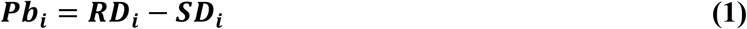

The perceptual bias of direction includes two components, systematic bias (*Sb*) and the nonsystematic bias (*NSb*_*i*_), as shown in Figure 1D. The systematic bias was calculated by the mean of perceptual biases of all directions:

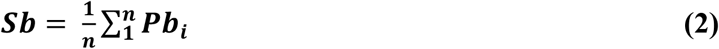

where n is eight in this study. Then a linear model was used to fit the systematic bias. The nonsystematic bias of direction was fitted by the cosine function with moment = 2 for generating two full oscillations in one complete direction cycle as follow:

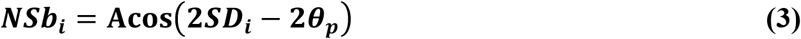

where the amplitude and phase of the peak was defined as A and *θ*_*p*_, respectively.

To evaluate tactile speed perception, continuous speed trajectories recorded from the touchpad were analyzed. Because substantial inter-subject variability was observed in baseline speed estimations, perceived speeds were normalized within each participant relative to the 80 mm/s reference condition. The relationship between normalized perceived speed and logarithm (base 10) of actual stimulus speed was then modeled using linear regression:

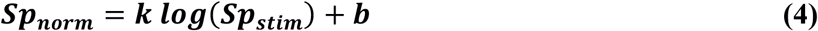

where the slope parameter *k* serves as a modulation index, representing the individual’s kinematic sensitivity to tactile speed variations. Larger absolute value of *k* indicates stronger modulation of perceived speed in response to changes in stimulator speed.

### 2.3 MRI acquisition and preprocessing

MRI data were acquired using 3.0T MRI system (PRISMA, Siemens Healthcare, Germany) equipped with a 20-channel head coil. T1-weighted anatomical images were obtained using a three-dimensional magnetization-prepared rapid gradient-echo imaging (3D-MPRAGE) sequence with the following parameters: repetition time (TR) = 2300 ms, echo time (TE) = 2.32 ms, flip angle = 8°, matrix size = 256×256, and field of view (FOV) = 240×240 mm^2^. Rs-functional images were acquired using a gradient echo-echo planar imaging (EPI) sequence with the following parameters: TR = 3000 ms, TE = 26 ms, matrix = 64×64, field of view = 228×228 mm^2^, flip angle = 90°, slice thickness = 3 mm, 48 axial slices, 120 volumes). During scanning, participants were instructed to remain awake with their eyes open while fixating on a central crosshair and minimizing movement or cognitive activity.

Rs-fMRI preprocessing was conducted using CONN Functional Connectivity Toolbox (version 22) (Whitfield-Gabrieli and Nieto-Castanon 2012) and SPM25 (https://www.fil.ion.ucl.ac.uk/spm/). The first 10 images were discarded to allow for signal stabilization. Remaining images underwent slice-timing correction and realignment for head motion correction. Participants exhibiting head motion exceeding 2 mm translation or 2° rotation were excluded from further analyses. Framewise displacement (FD) was calculated to quantify instantaneous head motion, and time points with FD > 0.5 mm were identified as motion outliers and scrubbed. To minimize physiological and motion-related confounds, nuisance regression was performed including linear trends, the 24-parameter Friston motion model, scrubbed volumes, and mean signals from the whole brain, white matter, and cerebrospinal fluid. Global signal regression (GSR) was additionally applied to enhance the specificity of functional connectivity estimates and reduce motion-related artifacts. The residual time series were then bandpass filtered (0.01–0.08 Hz).

For spatial normalization, individual T1-weighted image was co-registered to the mean functional image and segmented into gray matter, white matter, and cerebrospinal fluid. Functional images were normalized to the Montreal Neurological Institute (MNI) space using DARTEL and resampled to 3-mm isotropic voxels. Finally, normalized images were spatially smoothed using a Gaussian kernel of 6-mm full width at half maximum (FWHM) to improve signal-to-noise ratio and inter-subject alignment.

### 2.4 ROI-based functional connectivity analysis

To investigate network-level alterations underlying tactile-motion deficits in PD, we constructed a customized regions of interest (ROI)-based sensorimotor network encompassing cortical sensorimotor regions and cortico-basal ganglia-thalamo-cortical (CBGTC) circuits. A total of 26 bilateral ROIs were defined based on the HCPex atlas(Huang et al. 2022).

Cortical ROIs included primary regions directly mapped from the atlas, including Brodmann areas 4, 3a, 3b, 1, and 2. Higher-order associative regions were additionally defined by combining anatomically related HCPex parcels to generate composite representations of the supplementary motor area (SMA complex; 6ma, 6mp, SCEF, SFL), Area 5 (5L, 5m, 5mv), and Area 7 (7AL, 7Am, 7PC, 7PL, 7Pm, 7m). Subcortical ROIs included the caudate, putamen, external globus pallidus (GPe), internal globus pallidus (GPi), and thalamus. The thalamus ROI was defined by combining sensorimotor-related nuclei, including the ventral anterior (VA), ventral lateral anterior (VLa), ventral lateral posterior (VLp), ventral posterior lateral (VPL), parafascicular (Pf), central lateral (CL), and centromedian (CM) nuclei.

For ROI-to-ROI functional connectivity analysis, mean BOLD time series were extracted from each ROI, and pairwise Pearson correlation coefficients were computed between all ROI pairs. Correlation coefficients were subsequently transformed into Fisher z-scores prior to group-level statistical analyses.

### 2.5 Statistical Analysis

All statistical analyses were performed using python (version 3.11.13) with the *pingouin* and *scipy* packages. Demographic and clinical data were compared between groups using independent-samples *t*-tests for continuous data and Chi-square test for categorical data. Normality and homogeneity of variance were assessed using the Shapiro-Wilk test and Levene’s test, respectively.

For the tactile-motion perception experiments, directional bias and speed perception measures were analyzed using mixed-design analysis of variance (ANOVA), with Group (PD_AR_, HC) as a between-subject factor and Stimulus Speed (six levels) as a within-subject factor. Subject identity was included as a random factor to account for repeated measures. Post hoc pairwise *t*-tests with Bonferroni correction were conducted when appropriate. For psychophysical modulation analysis, the slope parameter (*k*) extracted from individual linear regressions was compared between groups using independent-samples *t*-tests.

For the resting-state functional connectivity analyses, group-level differences in functional connectivity were evaluated using a General Linear Model (GLM) with age and head motion included as covariates. Statistical significance was determined using network-level false discovery rate (FDR) correction at p < 0.05.

## 3 Results

### 3.1 Participant characteristics

Demographic and clinical characteristics of all participants are summarized in Table 1. No significant differences were observed between the PD_AR_ and HC groups in the age and sex distribution. Similarly, cognitive performance, including MMSE and MoCA, did not differ significantly between groups (both *p* > 0.1). Compared with HCs, PD_AR_ patients exhibited significantly higher scores on the Hamilton Depression Rating Scale (HAMD; *p* = 0.001) and Hamilton Anxiety Rating Scale (HAMA; *p* = 0.028), indicating mild affective symptoms. Most patients were classified as H-Y stage I-II, consistent with early-to-moderate disease severity. Mean framewise displacement during fMRI acquisition did not differ significantly between groups (*p* = 0.489), indicating comparable head motion during scanning.

**Table 1.** Demographic and clinical characteristics of participants.

| Data | HC ($n=21$) | PD<sub>AR</sub> ($n=25$) | $p$-values |
| --- | --- | --- | --- |
| Age (years) | 53.90 ± 1.08 | 57.12 ± 1.44 | 0.098<sup>a</sup> |
| Gender (F/M) | 12/9 | 13/12 | 0.959<sup>b</sup> |
| H-Y stage (I/II/III) | - | 10/14/1 | - |
| HAMD | 3.62 ± 0.61 | 7.48 ± 0.87 | 0.001<sup>a</sup> (*) |
| HAMA | 3.76 ± 0.65 | 5.92 ± 0.65 | 0.028<sup>a</sup> (*) |
| MMSE | 27.48 ± 0.42 | 26.16 ± 0.72 | 0.273<sup>a</sup> |
| MoCA | 25.10 ± 0.72 | 23.67 ± 0.77 | 0.192<sup>a</sup> |
| MDS-UPDRS II | - | 9.16 ± 0.55 | - |
| MDS-UPDRS III | - | 25.08 ± 2.49 | - |
| Mean FD (mm) | 0.082 ± 0.009 | 0.073 ± 0.009 | 0.489<sup>a</sup> |

Data is shown as mean ± standard error of the mean (SEM). H-Y stage, Hoehn and Yahr stage; MMSE, Mini-mental State Examination; MoCA, Montreal Cognitive Assessment; UPDRS, Unified Parkinson's Disease Rating Scale; FD, framewise displacement; -, not applicable; <sup>a</sup> two-sample t-test and <sup>b</sup> chi-squared test.

### 3.2 Impaired tactile-motion perception in PD_AR_

Psychophysical analyses revealed significant impairments in both tactile-motion direction and speed perception in PD_AR_ patients relative to HCs (Figure 2). For tactile-motion direction perception, PD_AR_ participants exhibited significantly larger systematic bias compared with HCs (PD = -14.66°±2.70°,HC = -2.86°±4.92°; t(46)=2.11, *p* = 0.047). Analysis of the non-systematic bias revealed no significant group difference in the modulation amplitude (t(46)=1.01, *p* = 0.324), whereas the phase shift differed significantly between two groups (t(46)=2.53, *p* = 0.021). These results indicate altered spatial representation of tactile-motion direction in PD_AR_ participants (Figure 2C).

**Figure 2.**
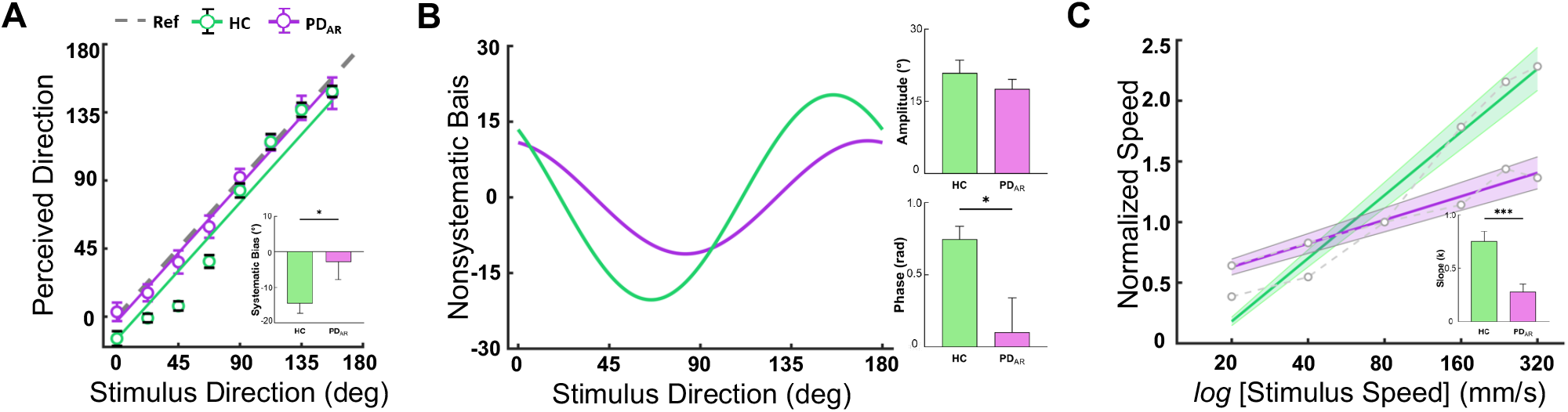
Psychophysical Results. (A) Perceived direction. Grey dotted line was the zero-bias as the reference line. (B) Cosine fitting curves for nonsystematic bias in HC (green) and PD_AR_ (purple) groups. (C) Linear fitting for normalized speed of HC (green) and PD_AR_ (purple) groups. Gray dots represent the mean perceived speed within each group. Independent-samples t-tests were performed for systematic bias, nonsystematic bias (amplitude and phase), and the slope (k) of perceived speed. \**p* < 0.05, \*\*\**p* < 0.001.

For tactile-motion speed perception, a mixed-design ANOVA with Group (HC, PD) and Speed Condition (20, 40, 80, 160, 240, and 320 mm/s) revealed a significant main effect of Speed (F(1,28)=69.83, *p* < 0.0001), confirming that perceptual responses systematically increased with actual stimulus speed. Although the main effect of Group did not reach statistical significance (F(1,28)=3.58, *p* = 0.069), a significant Group × Speed interaction was observed (F(1,28)=5.39, *p* = 0.028). Post-hoc pairwise comparisons with Benjamini-Hochberg FDR correction revealed that the perceptual deficits in PD were speed-dependent. Specifically, PD participants exhibited significantly reduced perceived speed at mid-to-high stimulus speeds (Speeds 4–6; all *q* = 0.0405), whereas their performance was comparable to that of HCs at lower speeds (Speeds 1–3; all *q* > 0.7).

Crucially, to quantify this perceptual blunting across the speed spectrum, we compared the psychophysical modulation index (slope parameter *k*) extracted from the linear regression for each participant. Consistent with the ANOVA results, an independent t-test demonstrated a significantly flattened perceptual slope in the PD group compared to HCs (mean Slope: PD = 0.28 vs. HC = 0.75; t(46)=3.99, *p* < 0.001). This marked reduction in the slope parameter further indicates that PD_AR_ patients exhibit substantially diminished kinematic sensitivity to dynamic changes in tactile-motion speed perception.

### 3.3 Cortical hypoconnectivity and subcortical hyperconnectivity in PD_AR_

Resting-state functional connectivity analyses revealed distinct patterns of cortical hypoconnectivity and subcortical hyperconnectivity in PD_AR_ patients relative to healthy controls (Figure 3A and Table 2). Within the cortical somatosensory network, PD_AR_ patients exhibited significantly reduced functional connectivity among primary somatosensory regions, including Brodmann Areas 3a, 3b, 1, and 2, as well as between higher-order somatosensory association regions encompassing Areas 5 and 7 (all *p*-FDR < 0.05). The most prominent reductions were observed between bilateral Area 1 regions and between contralateral Area 3b and ipsilateral primary motor cortex (M1), indicating reduced functional connectivity within and between sensorimotor regions. Consistently, the mean intra-somatosensory functional connectivity strength was significantly lower in the PD_AR_ group compared with HCs (t(44) = 2.67, *p* = 0.011). In contrast, subcortical-cortical connectivity analyses demonstrated significantly increased coupling between basal ganglia-thalamic regions and cortical sensorimotor areas in PD_AR_ patients. Specifically, enhanced connectivity was observed between the thalamus and M1, as well as between the globus pallidus internus (GPi) and both M1 and S1 regions (all *p*-FDR < 0.05). These increases were predominantly localized to the hemisphere corresponding to the clinically more-affected side. Together, these findings demonstrate a network-level imbalance characterized by reduced cortical somatosensory integration alongside enhanced subcortical-cortical coupling in PD_AR_ patients.

**Table 2.**
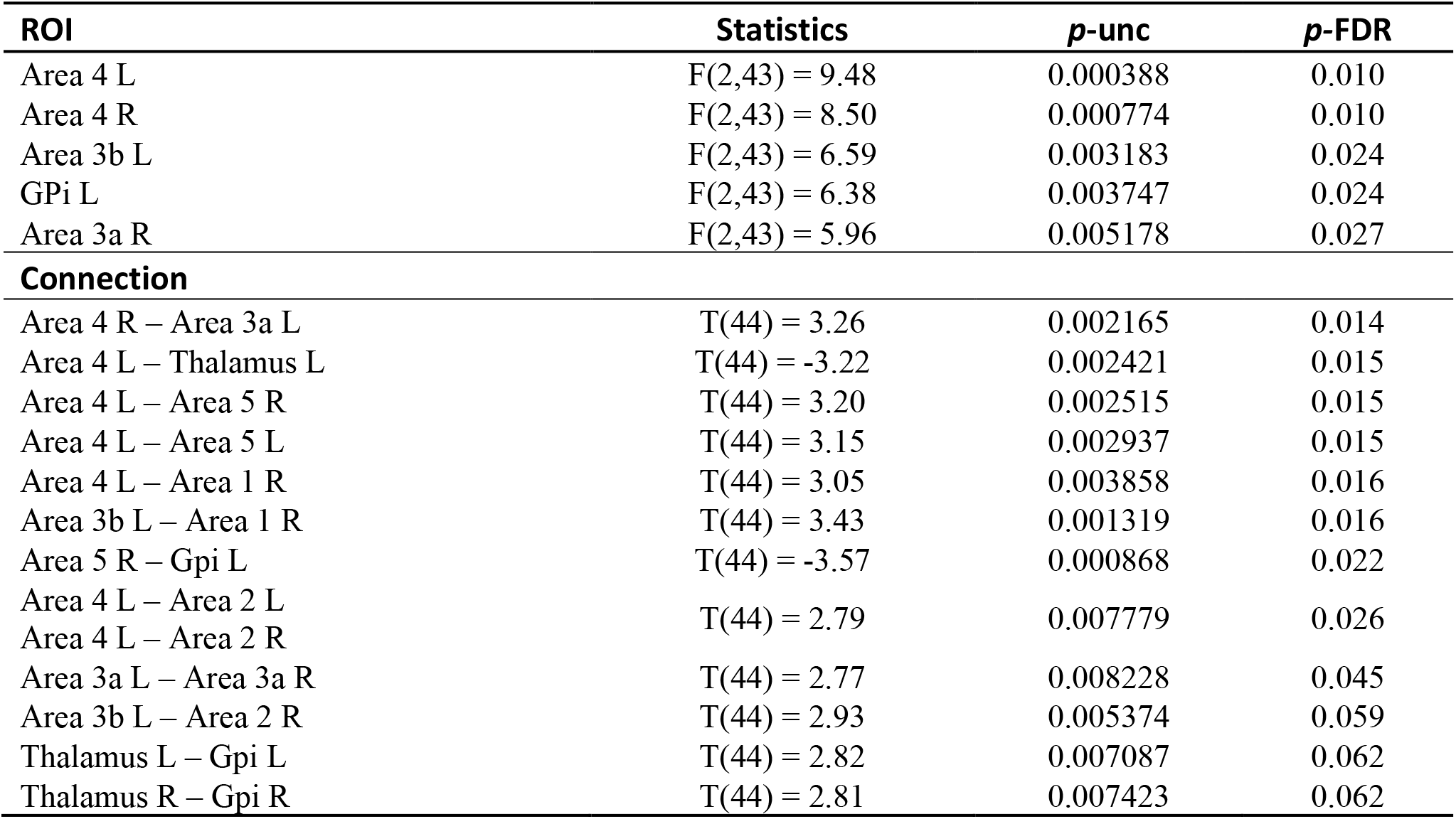
Statistics Results for Functional Connectivity Changes between HC -PD.

**Figure 3.**
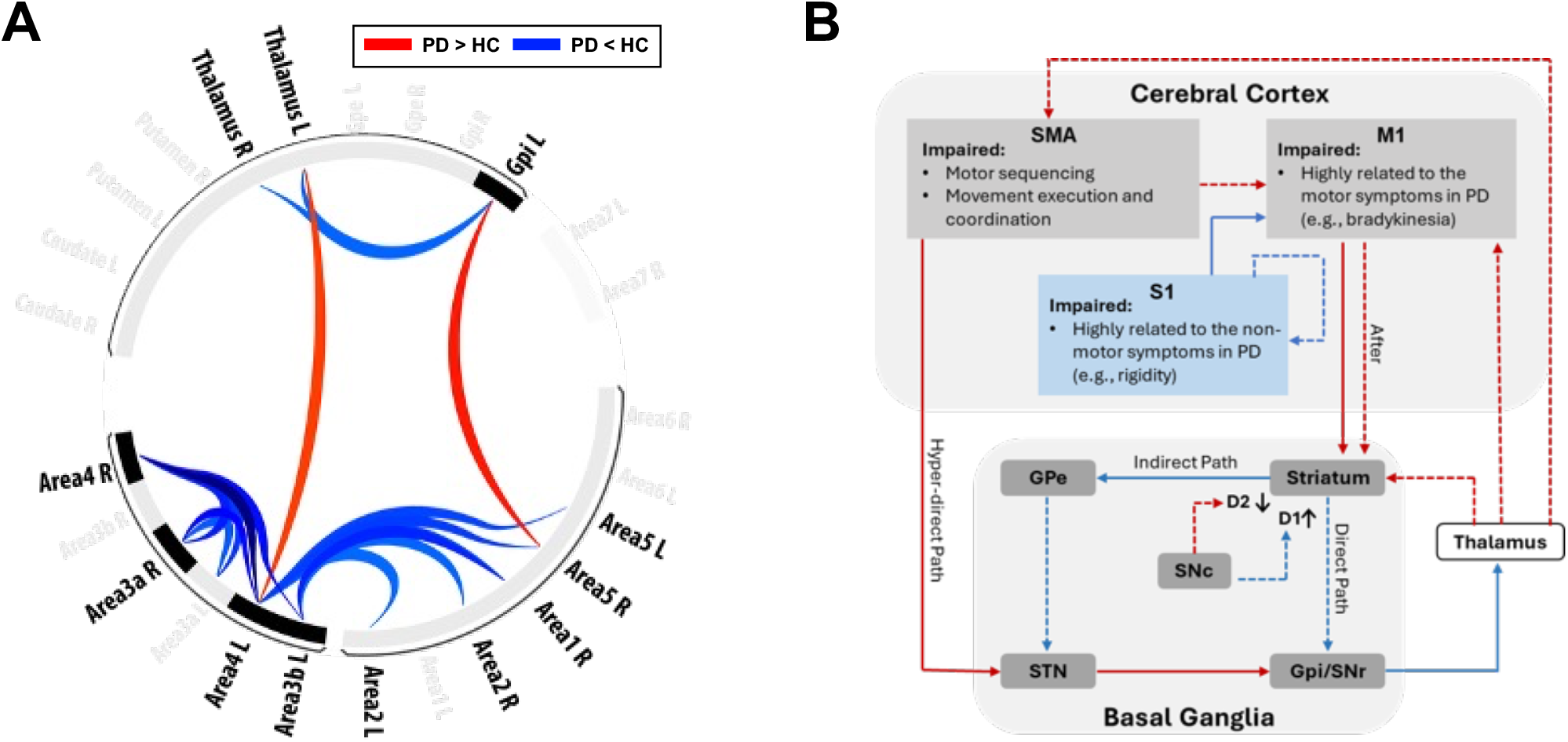
Functional Connectivity Alterations and Cortico–Basal Ganglia–Thalamo–Cortical Network Reorganization in PD_AR_. (A) Group differences in resting-state functional connectivity between PD_AR_ patients and healthy controls (HCs). Blue lines represent significantly reduced functional connectivity in PD_AR_ relative to HCs, whereas red lines indicate significantly increased functional connectivity. Statistical significance for both nodal properties (F-tests) and functional edges (t-tests) was set at *p* < 0.05, FDR-corrected. (B) Schematic illustration of altered cortico-basal ganglia-thalamo-cortical (CBGTC) network organization in PD_AR_. Based on the classical Parkinsonian circuit model, the schematic incorporates the present findings highlighting disrupted somatosensory cortical integration, particularly involving the primary somatosensory cortex (S1), and its altered functional interactions with the primary motor cortex (M1) and supplementary motor area (SMA).

## 4 Discussion

The present study combined psychophysical assessment of tactile-motion perception with resting-state fMRI to investigate somatosensory dysfunction in akinetic-rigid PD. PD_AR_ patients exhibited significant impairment in both directional and speed-dependent tactile-motion perception, accompanied by widespread reductions in cortical somatosensory connectivity and increased subcortical-cortical coupling within the CBGTC network. These findings suggest that tactile dysfunction in PD may arise from a network-level imbalance characterized by disrupted cortical somatosensory integration and abnormal subcortical-cortical interactions.

### 4.1 Impaired spatiotemporal processing of tactile-motion in PD_AR_

The present findings extend previous studies of somatosensory dysfunction in PD_AR_ by demonstrating abnormalities not only in basic tactile perception, but also in higher-order tactile-motion processing. Previous work has primarily focused on tactile detection thresholds, vibration discrimination, or proprioceptive impairments(A. Conte et al. 2013; Antonella Conte et al. 2017; Kesayan et al. 2015; Konczak et al. 2012; Lee et al. 2018), whereas the current study examined the perception of dynamically moving tactile stimuli. Because tactile-motion perception depends on the integration of spatial and temporal sensory information(Patel et al. 2014), the observed impairments suggest that somatosensory dysfunction in PD extends beyond elementary sensory deficits.

PD_AR_ patients exhibited a pronounced systematic directional bias during tactile-motion perception, whereas the nonsystematic component of their responses remained relatively preserved. Specifically, the amplitude of trial-by-trial variability did not differ significantly between groups, yet the phase of the fitted cosine function was significantly shifted. This pattern demonstrates that PD does not simply increase perceptual noise but instead alters the spatial transformation of tactile-motion information. Such directional displacement may reflect abnormalities in transforming tactile signals from skin-centered coordinates into external spatial reference frames (Badde and Heed 2023; Chen et al. 2020; Heed et al. 2016; Qu et al. 2025).

Speed perception revealed a complementary but distinct pattern of impairment. While PD_AR_ patients performed comparably to healthy controls at low stimulus speeds, perceived speed was significantly reduced at intermediate to high-speed stimulation. The flattened psychophysical modulation slope further indicates diminished sensitivity to dynamic changes in tactile-motion speed. Importantly, this speed-dependent impairment suggests that tactile-motion deficits in PD become more pronounced under conditions requiring rapid temporal integration of continuously evolving sensory input (Merchant et al. 2013). These findings suggest that tactile-motion deficits in PD become particularly pronounced when temporal processing demands exceed the capacity of compromised sensorimotor circuits (A. Conte et al. 2013).

These findings are consistent with previous evidence demonstrating abnormal temporal discrimination and impaired sensorimotor integration in PD (Artieda et al. 1992; Antonella Conte et al. 2017; Lee et al. 2018). Perceiving speed requires precise coordination of sensory population responses and relies on intact cortical timing mechanisms (Voudouris and Fiehler 2021). Given that PD is associated with abnormal basal ganglia output and disrupted cortical synchrony, the reduced ability to track high-speed tactile-motion may reflect impaired neural timing and abnormal temporal coordination within sensorimotor circuits (Avanzino et al. 2016; Hammond et al. 2007). The partial dissociation between direction and speed impairments further suggests that PD disrupts multiple functionally distinct components of tactile-motion processing, including a spatial reference-frame mechanism governing direction perception and a temporally precise integration mechanism governing speed perception (Y.-C. Pei et al. 2011). Together, the present behavioral findings suggest that tactile-motion dysfunction in PD_AR_ patients involves impairments in both spatial representation and temporal integration.

### 4.2 Disrupted somatosensory integration in PD_AR_

Resting-state functional connectivity analyses revealed widespread reductions in intrinsic connectivity within cortical somatosensory regions in PD_AR_ patients, involving both primary somatosensory regions (Area 3a, 3b, 1, and 2) and higher-order somatosensory association regions (Area 5 and 7). These affected regions span multiple hierarchical stages of tactile processing(Abbruzzese and Berardelli 2003; Avanzino et al. 2016; Ryun et al. 2023), suggesting impaired intrinsic coordination within the cortical somatosensory network. Among these alterations, reduced connectivity between S1 and M1 was particularly prominent. Functional interactions between somatosensory and motor cortices are essential for transforming tactile inputs into sensorimotor representations that support perception and action (Ebrahimi and Ostry 2024). Accordingly, weakened S1-M1 connectivity may contribute to the altered directional perception patterns observed behaviorally. Importantly, the preservation of nonsystematic variability alongside altered phase parameters suggests that lower-level sensory encoding may remain relatively intact, whereas higher-order spatial organization processes become disrupted. This dissociation is consistent with previous reports of impaired sensorimotor integration in PD (Abbruzzese and Berardelli 2003; Y. Wang et al. 2024).

Reduced intra-somatosensory connectivity may also contribute to impaired temporal integration during tactile-motion processing. Accurate perception of moving tactile stimuli requires coordinated cortical activity for encoding motion direction, speed, and spatiotemporal continuity(Y.-C. Pei et al. 2011; Y. C. Pei and Bensmaia 2014). Collectively, these findings extend previous rs-fMRI studies of sensorimotor dysfunction in PD(Goelman et al. 2021; Permezel et al. 2023; Shen et al. 2024; S. Wang et al. 2021; Y. Wang et al. 2024) and suggest that disrupted cortical network organization contributes specifically to impairments in higher-order tactile-motion processing.

### 4.3 Subcortical hyperconnectivity and network reorganization

In contrast to the reduced connectivity observed within cortical somatosensory regions, PD_AR_ patients exhibited increased functional coupling between basal ganglia-thalamic structures and cortical sensorimotor areas, particularly involving the thalamus, GPi, M1, and S1. These findings indicate that PD is associated with altered subcortical-cortical network organization within the CBGTC system. Abnormal synchronization and altered functional coupling within basal ganglia-thalamocortical circuits have been consistently reported in PD [31,37–40]. Increased connectivity involving the thalamus and GPi may reflect altered information flow within sensorimotor pathways secondary to dopaminergic dysfunction. Although the precise functional significance of this hyperconnectivity remains unclear, increased subcortical-cortical coupling may alter the precision of cortical sensory processing by disrupting the balance of network-level excitatory and inhibitory interactions. Interestingly, increased subcortical connectivity coexisted with reduced cortical somatosensory integration. This dissociation suggests that cortical and subcortical networks may undergo distinct forms of reorganization in PD. While cortical somatosensory regions exhibited reduced intrinsic coordination, basal ganglia-thalamic circuits showed enhanced connectivity with sensorimotor cortical areas. Together, these findings support the view that tactile dysfunction in PD emerges from distributed sensorimotor network abnormalities rather than isolated dysfunction within a single cortical region. The present findings may also help explain why tactile impairments in PD frequently coexist with motor dysfunction. The basal ganglia-thalamocortical system plays a central role in integrating sensory information with motor planning, execution and cognitive processes (Baudrexel et al. 2011; Roth and Ding 2024; Zhuang et al. 2023). Accordingly, abnormal subcortical-cortical interactions may contribute both to impaired tactile-motion perception and broader sensorimotor dysfunction in PD.

### 4.4 Integrated network perspective of tactile dysfunction in PD_AR_

By combining psychophysical assessment with resting-state functional connectivity analysis, present study demonstrates that tactile-motion perceptual deficits in PD coexist with large-scale alterations in sensorimotor network organization. Specifically, impairments in directional and speed-dependent tactile-motion perception coexisted with reduced cortical somatosensory connectivity and increased subcortical-cortical coupling. These findings support a network-level framework in which disrupted cortical sensory integration impairs higher-order spatiotemporal tactile processing, while altered basal ganglia-thalamocortical connectivity reflects broader sensorimotor network reorganization. Importantly, the present findings suggest that somatosensory dysfunction in PD should not be viewed solely as a secondary consequence of motor impairment, but rather as part of a distributed disturbance involving both sensory and motor systems (Patel et al. 2014).

### 4.5 Limitations and future directions

Several limitations should be acknowledged. First, the sample size was relatively modest, particularly for the psychophysical experiments, which may limit statistical power and generalizability, and the stability of subgroup-specific network findings. Second, the cross-sectional design precludes causal inference regarding the relationship between altered functional connectivity and tactile perceptual deficits. Third, although behavioral impairments and network abnormalities were observed concurrently, direct brain–behavior correlation analyses were not performed. Future studies incorporating larger cohorts, longitudinal designs, and multimodal approaches may help clarify how alterations in somatosensory network organization contribute to the progression of tactile dysfunction in PD. In addition, only akinetic-rigid dominant PD patients were included in the present study. Because distinct PD subtypes may exhibit different sensorimotor network alterations, future work should compare tactile-motion processing across clinical subtypes. Combining functional neuroimaging with electrophysiological or neuromodulation approaches may further clarify the neural mechanisms underlying tactile-motion dysfunction and help identify potential therapeutic targets for restoring sensorimotor integration in PD.

## 5 Conclusion

In summary, the present study demonstrates that tactile-motion deficits in Parkinson’s disease are accompanied by distinct alterations in sensorimotor network organization, characterized by reduced cortical somatosensory connectivity and increased subcortical-cortical coupling. These findings suggest that higher-order tactile dysfunction in PD arises from disrupted spatiotemporal integration within distributed sensorimotor networks rather than isolated sensory impairment alone. By linking tactile-motion perceptual abnormalities with intrinsic network alterations, this study provides new insight into the neural basis of somatosensory dysfunction in PD and highlights the somatosensory system as a potential target for future neuromodulatory and sensory-based therapeutic strategies.

## 6 Data availability statement

The datasets generated and analyzed during the current study are available from the corresponding author upon reasonable request.

## 7 Conflict of interest

*The authors declare that the research was conducted in the absence of any commercial or financial relationships that could be construed as a potential conflict of interest*.

## 8 Generative AI statement

The authors declare that no Gen AI was used in the creation of this manuscript.

## 9 Author contributions

B. Qu: Writing – original draft, Methodology, Formal analysis, Visualization; Y. Jiang: Writing – original draft, Investigation; Y. Yue: Investigation; T. Shen: Investigation; C. Gao: Data curation, Formal analysis; D. Huang: Formal analysis; Y. Shen: Writing – review & editing, Funding acquisition; C. D. Zeeuw: Writing–review & editing; B. Zhang: Resources, Supervision; H. Lai: Conceptualization, Writing–review & editing, Supervision, Funding acquisition.

## 10 Funding information

This work was supported by grants from Key Strategic Science and Technology Cooperation Project (2023YFE0206800, to Y. S.) of the Ministry of Science and Technology of China and the National Natural Sciences Foundation of China (81600982 and 82541160282, to HY. L.).

## Notes

### Competing Interest Statement

The authors have declared no competing interest.

